# Variation in diet and microbial exposure shape the performance of the Asian tiger mosquito, *Aedes albopictus*

**DOI:** 10.1101/2022.11.16.516852

**Authors:** Vincent Raquin, Edwige Martin, Guillaume Minard, Claire Valiente Moro

## Abstract

Along their life cycle, mosquitoes colonize different ecological niches with various microorganisms and diet sources that likely modulate their performance *(i.e.* a set of mosquito fitness-related traits). However, which ecological parameters and how their variations modulate mosquito performance is not completely understood. In this study, we used *Ae. albopictus* surface-sterilized eggs re-associated or not to conventional bacterial microbiota upon a range of diet concentrations and addressed the impact of microbial inoculum and diet concentration variation on several mosquito performance traits. Results showed that mosquito juvenile survival depends on the interaction between bacterial inoculum load and diet concentration in the breeding water. Exposure to bacteria in rearing water shorten larval development time although it impacted larvae survival in an inoculum and diet concentration-dependent manner. Bacterial composition of larval rearing water was mainly structured by the bacterial inoculum concentration, with some Operational Taxonomic Units abundances correlating with larval traits. *Ae. albopictus* survival, development and bacterial community patterns upon gradients of diet and bacterial inoculum illustrated the complex impact of diet-microbiota interaction on mosquito performance. These findings argue the importance of deciphering host-microbe-environment interactions and open promising perspectives to improve *Ae. albopictus* control measures in the field.

**IMPORTANCE:** Microbiota is increasingly recognized as a driving force of metazoan biology, impacting diverse traits including nutrition, behaviour or reproduction. The microbial impact on host nutrition is among the most studied host-microbe interactions although it remains poorly understood in arthropod vectors like mosquitoes. Here, we manipulated mosquito microbiota using gnotobiology to decipher the impact of bacteria and diet on the Asian tiger mosquito, *Ae. albopictus.* These results are key to understand the link between diet and bacteria concentration on juvenile mosquitoes as well as carry-over effects in adults. They unveil some specific aspects of mosquito-bacteria interactions while opening interesting avenues for vector management of this vector of arboviruses.

## INTRODUCTION

From mammals to insects, host-microbiota interactions drive host nutrition, behavior, reproduction or development thereby impacting host fitness (1,2). However, the ecological determinants governing these symbiotic interactions remain unknown in many systems. During nutritional symbioses, microbial symbionts benefit to host by serving as nutrient source, detoxifying diet, participating in nutrient digestion/intake or supplementing diet with essential molecules (2, 3). Converging evidences show that diet is a driving force of symbiosis, for instance by selecting for diet-adapted microbial communities (4, 5). Advances in the field of symbiotic interactions benefited from the development of gnotobiology *(i.e.* study of hosts with a controlled microbiota) as well as holidic diets *(i.e.* synthetic nutrient source made of pure chemical components). In insects, individuals are deprived from environmentally-acquired microorganisms by surface sterilization of eggs that are put to hatch under sterile environment (6). Individuals can then be re-associated to selected microorganisms to interrogate microbial influence on host traits, under varying (a)biotic conditions such as diet. However, holidic diets and gnotobiotic models are unevenly available among host species (8) although attempts are made to develop gnotobiology and holidic diets in mosquitoes (9, 10).

Compelling evidences from the fruit fly *Drosophila melanogaster* obtained by comparing axenic with gnotobiotic individuals support that bacteria are not only a source of nutrients but establish a diet-dependent nutritional symbiosis to provide their host with essential factors when raised on scarce diets (11–13). However, in mosquitoes the impact of the microbiota on mosquito performance under the influence of the diet in a still poorly understood way. Mosquitoes *(Diptera: Culicidae)* are holometabolous insects with four distinct life stages (egg, larvae, pupae and adult). Larval and pupal stages develop in aquatic breeding sites before winged adults emerge. The breeding site provides larvae with nutrients to complete their development (14–16) and shapes their microbiota (17–19). Diet quantity and composition impact mosquito larval growth, development time and survival (20–25) as well as adult immunity, vector competence and fitness (26–30). Natural larval water habitats contain variable resource inputs that stimulate microbial growth, with the decomposition of detritus by microorganisms releasing nutrients uptake by larvae (31–33). In turn, microbiota also deeply impacts mosquito biology, from larval to adult stage (18). Supplementation of sterile larval rearing water with living bacteria seemed mandatory to reach adult stage (34, 35). However, optimal diet and rearing conditions can lead to complete larval development in absence of environmental microorganisms, as demonstrated in a proof-of-concept study in *Ae. aegypti* (36). Similarly to what was reported in *D. melanogaster* (37, 38) axenic mosquito larvae present major physiological changes including larval developmental delay, reduced adult size or extended adult life span that correlate with specific transcriptomic profiles compare to conventional larvae (36, 39, 40). More recently, study of axenic *Ae. aegypti* showed that bacteria in rearing water mediate larval nutrient sensing and growth activation by provisioning larvae with essential vitamins (41, 42). In *Ae. aegypti,* bacteria-mediated growth-promoting effect on larvae depends on both the bacterial strain and diet type (rat chow or fish food) (43). Microbial supplementation of water as unique nutrient source is not as efficient as synthetic rich diet in promoting larval survival and developmental rate in conventionally reared *(i.e.* non axenic) individuals, with major variations in larval performance being observed according to the microbial strains used (44). This underlines that an interaction between microbiota and diet during larval stage impacts performance in *Ae. aegypti* larvae, with potential carry-over effects on adult traits (26, 45). Recent data in *Ae. aegypti* suggest that diet *x* microbiota interaction occurs as diet concentration impacts bacterial abundance and composition in rearing water and larvae (46). However, additional data are needed to assess if diet and microbiota have similar effects on the biology of other mosquito species, and if and how diet x microbiota interaction shapes mosquito performance.

The Asian tiger mosquito *Aedes albopictus* is an important vector of human pathogens including dengue virus, chikungunya virus or Zika virus (47, 48). This mosquito species thrives notably in urban and suburban environments where females lay eggs in a broad range of breeding sites (51). Field-collected specimens harbor a core bacterial microbiota although variations in bacterial diversity and community structure are observed according to geographic origin, developmental stage or sex (52). Bacteria impact host traits in *Ae. albopictus* including oviposition site selection (53), sugar feeding (54) or larval development (55). *Ae. albopictus* juvenile development also depends on diet as larvae present a developmental delay below a given diet concentration that microbiota, including the native intracellular bacterium *Wolbachia* cannot counterbalance (56). Together, it suggests that microbiota and diet impact *Ae. albopictus* performance. But to the best of our knowledge, no work generated axenic *Ae. albopictus* nor study the concomitant impact of diet and microbiota concentration on mosquito performance. *Ae. albopictus* larvae were successfully deprived from environmental microorganisms, with the exception of intracellular ones such as *Wolbachia*, while allowing larval development up to the adult stage. In order to disentangle the relative and combined importance of both microbiota and diet on mosquito traits, the performance of gnotobiotic *Ae. albopictus* was investigated along a gradient of nutrient concentrations. Water microbial community composition and relative abundance were also determined across diet and inoculum conditions.

## RESULTS

### Altered microbiota larvae exhibit a diet-dependent juvenile developmental pattern compared to conventional siblings upon constant bacterial inoculum concentration

*Ae. albopictus* larvae naturally hosting the intracellular bacterium *Wolbachia* but deprived from environmental microorganisms were generated by egg surface sterilization (FIG S1), hereafter referred as altered microbiota (AM) larvae. A very low (about 8,600 fold less as estimated by 16S qPCR) residual amount of bacterial DNA in AM rearing water was still detected compare to conventional (CONV) water after egg sterilization (FIG S3). It was composed of bacterial OTUs either poorly abundant *(Aeromonas, Dysgonomonas, Pseudomonas)* and/or only found in highly diluted inoculum condition *(Bacteroides, Stenotrophomonas)* of CONV water samples (FIG S3 and FIG 4). Of note, AM larvae were stalled at larval stage only reached adult stage when incubated in darkness (data not shown) recapitulating axenic mosquito larvae development pattern from previous studies (36, 41).

The impact of diet concentration in rearing water on larval development was estimated using AM larvae or siblings re-associated upon hatching with a conventional microbial inoculum (CONV) composed of 5 to 6 major bacterial species (FIG S2). AM and CONV larvae were exposed to a gradient of diet from 0.1% to 20% concentration (FIG S4). Diet concentration modulates larvae development depending on the microbial status (Wald X^2^, *P*_diet x microbial status_ = 3.97e^−05^). Overall, CONV larvae developed from 12% to 0.5% diet concentration whereas AM larvae developed from 20% to 5% although larval viability reached higher values in AM compare to CONV (FIG S4). Indeed, when exposed to a single, low dilution (10^−3^) of inoculum, the proportion of CONV larvae that reached pupal stage remained below 25% regardless of diet concentration. In addition, an increase in water turbidity was noticed the day after inoculation with larval death occurring prior fourth instar.

To disentangle the impact of microbiota and diet concentration on mosquito larvae development, a set of three experiments was conducted in which larvae were exposed to a single dose of bacterial inoculum, but more diluted (10^−6^) to limit larval mortality while randomizing microbiota variation by loading a different bacterial inoculum in each experiment. Four diet concentrations spanning the viable range for AM and CONV larvae (2, 5, 10 and 12%) were tested. Results showed that larval development differs according to experiments (FIG 1), maybe due to variations in egg batch or composition of microbial inoculum (FIG S2). When controlling for random experiment effect, larvae-to-pupae viability depended on the interaction between microbial status and diet concentration (Wald X^2^, *P*_diet x microbial status_ < 2.2e^−16^) (FIG 1A). AM larvae survival from 5 to 12% diet concentration was similar but lower than at 2% whereas CONV larvae viability at 2 and 5% diet concentration was similar but higher than at 10 and 12% (FIG 1A). Unlike larvae, pupae-to-adult viability was not impacted by these two variables (Wald X^2^, *P*_diet_ = 0.16, *P*_microbial status_ = 0.31, *P*_diet x microbial status_ = 0.51) (FIG 1 B). Larval viability was higher in AM compared to CONV except at the lowest diet concentration of 2% (FIG 1A). Among the larvae that reached pupal stage, we measured the time (in days) needed to reach 50% of the final number of pupae (Day_50_) as a proxy of development time. As observed for larvae-to-pupae viability, mosquito juvenile development time depended on the interaction between microbial status and diet concentration (Anova, *P*_diet x microbial status_ < 2.2e^−16^) (FIG 1C). Overall, CONV larvae always developed faster (about 7 days) than AM counterparts. The Day_50_ of CONV larvae at 2, 5 and 10% diet concentrations were similar but smaller than at 12%, whereas the Day_50_ of AM larvae at 2% diet concentration was higher than at 5%, which in turn was higher than at 10 and 12% (FIG 1C).

**FIG 1.**
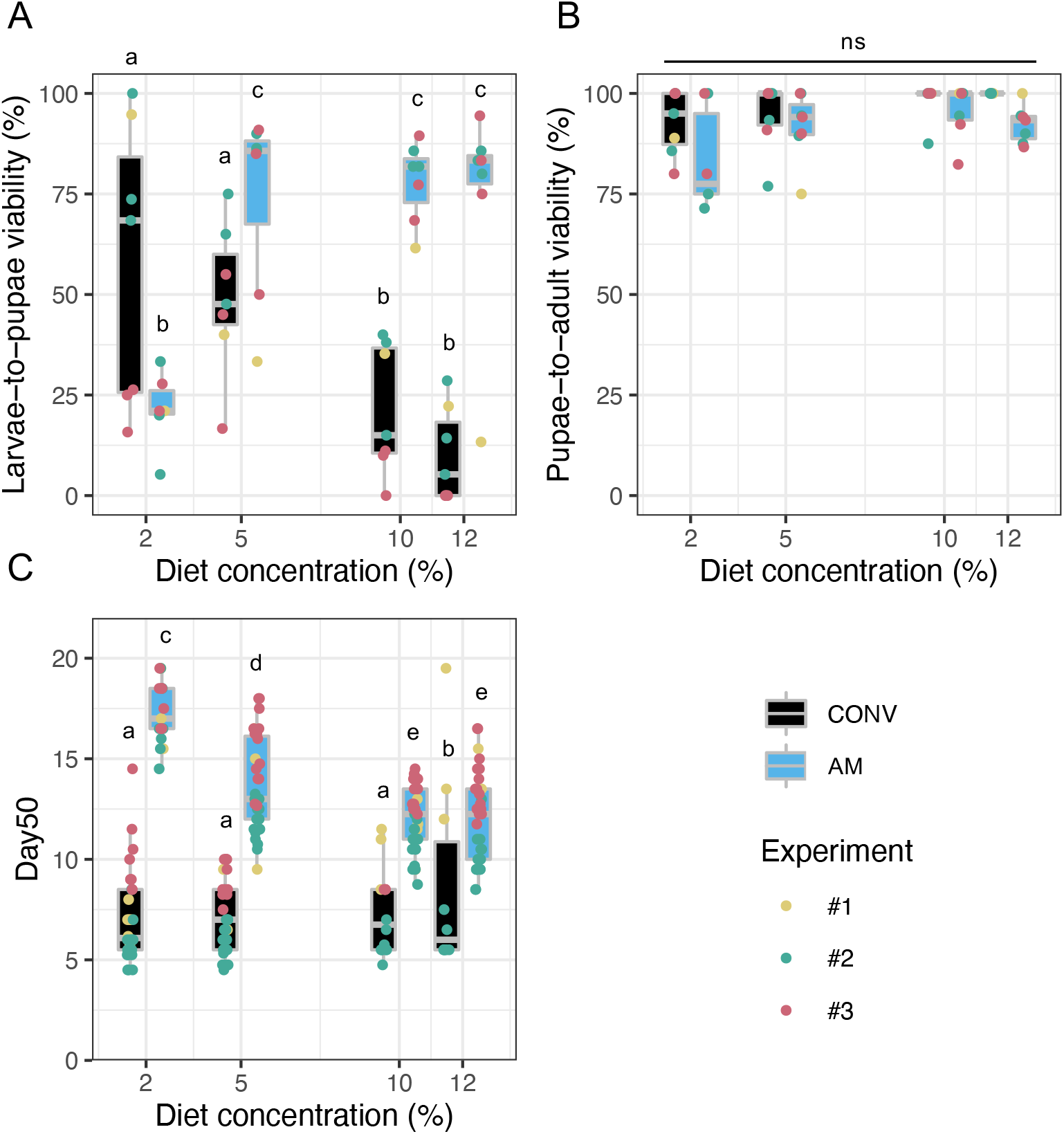
Juvenile development pattern of *Ae. albopictus* larvae upon a diet gradient in presence or absence of bacteria. Development pattern of CONV (black) and AM (blue) larvae at four diet concentrations (2, 5, 10 and 12%). Three independent experiments (#1 to #3 as indicated by color code) were performed, with one to three 6-well plates per experiment (3 larvae per well) for each microbial status and diet concentration combination. (A) Larval viability expressed as the proportion (in %) of mosquito larvae that reached pupal stage. Each dot represents the mean viability per 6-well plate. (B) Pupal viability expressed as the proportion (in %) of mosquito pupae that reached adult stage. Each dot represents the mean viability per 6-well plate. (C) Larval development time into pupae is represented as the time (in days) needed to reach 50% of the final number of pupae (Day_50_). Different letters indicate statistically significant viability following Tukey-HSD *post-hoc* pairwise comparison.

No significant difference was observed in the proportion of adults from each sex according to microbial status, diet concentration or their interaction (Wald X^2^, *P*_diet_ = 0.72, *P*_microbial status_ = 0.50, *P*_diet x microbial status_ = 0.067) (FIG 2A). Adult wing length was significantly impacted by the interaction between microbial status and diet concentration (Anova, *P*_diet x microbial status_ = 0.009) (FIG 2B). CONV females presented larger wings compare to males, and within each sex, wing length was not different according to diet concentration. In AM conditions, females displayed higher wing length compared to males except at 2% diet concentration, with all males presenting similar wing length regardless of diet concentration. Within AM females, a similar wing length was measured at 10 and 12%. However, AM females were larger at 12%, compared to 5 and 2% diet concentrations.

**FIG 2.**
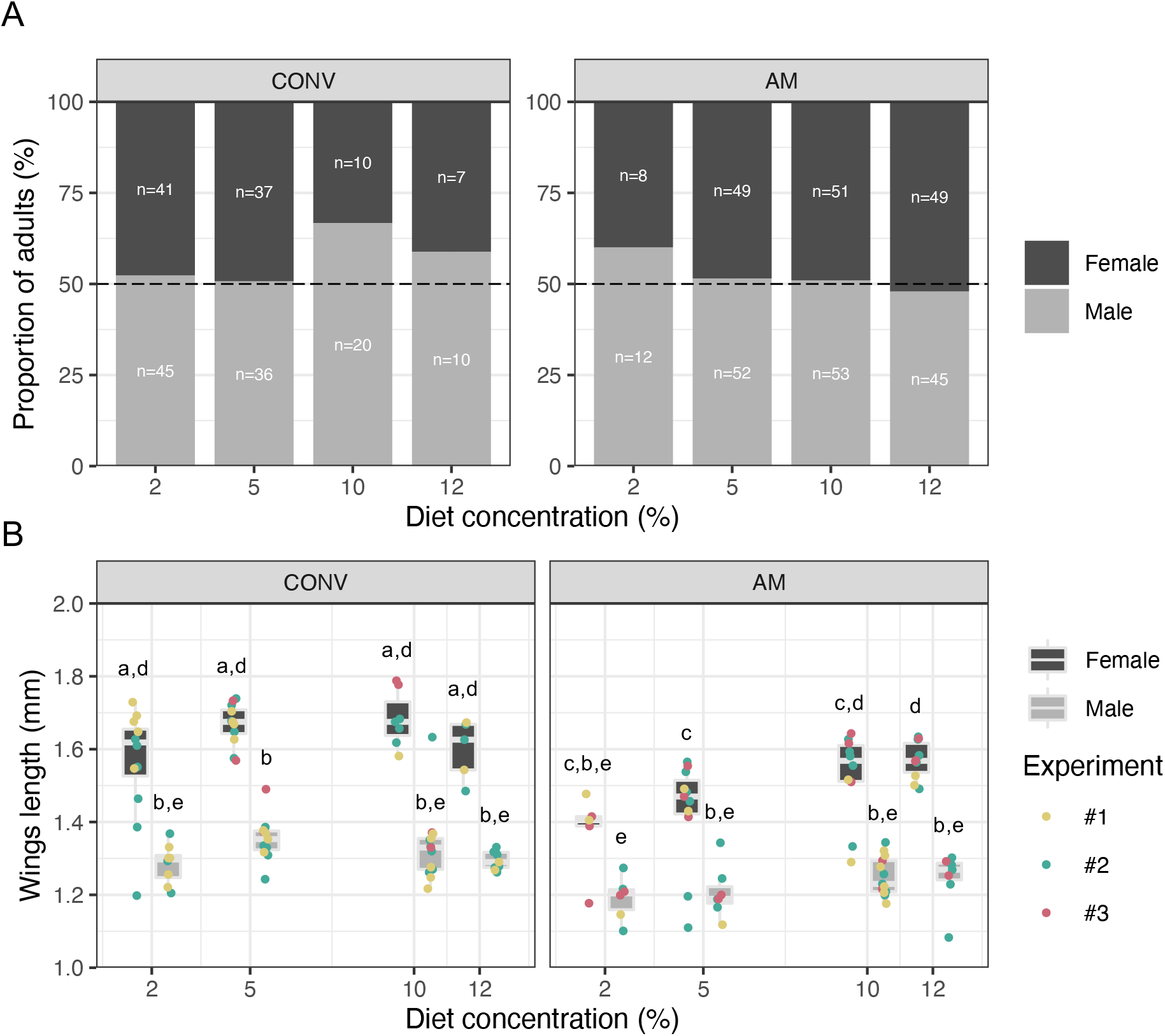
Adult sex ratio and wing size upon a diet gradient in presence or absence of bacteria. Adults derived from CONV or AM larvae reared at different diet concentrations (2, 5, 10 and 12%) were sorted by sex and individual wings length was measured. Three independent experiments (#1 to #3) were performed, with one to three 6-well plates per experiment (3 larvae per well) for each microbial status and diet concentration combination. The number of adults available varied according to the juvenile mortality rate of each combination. (**A**) Proportion of female and male adult mosquitoes. The number of individuals (n) for each condition is indicated. (B) Adult wings length (in mm). Each dot represents the mean length of both wings for an individual. From 5 to 17 individuals were analysed depending on the sex, microbial status and diet concentration. Different letters indicate statistically significant viability following Tukey-HSD *post-hoc* pairwise comparison.

### Bacterial load and diet concentration in larval rearing water are strong predictors of *Ae. albopictus* juvenile performance

We showed that at constant initial load, bacteria promote larval performance depending on diet concentration. In nature, microbial load and diet concentration are likely to vary in larval breeding sites. Therefore, we asked if and how concurrent variation in bacterial inoculum and diet concentration could shape mosquito juvenile performance. Within a single experiment, CONV larvae were exposed to three different dilutions of inoculum (10^−4^, 10^−6^ and 10^−8^) and three diet concentrations (1, 5 and 12%) prior to measure larval viability, development time and microbiota composition. Concentrations of inoculum and diet were chosen according to previous conditions showing the most contrasted effects on larval performance (FIG 1). Three independent batches of inoculum (B1, B2 and B3) were used. Analysis of the bacterial community variance of the water of each batch showed differences in bacterial composition (FIG S2). When controlling for inoculum batch variation, results showed that larval viability is impacted by the interaction between inoculum and diet concentration (Wald X^2^, *P*_diet x inoculum_ = 6.86e^−7^) (FIG 3A). At 10^−4^ inoculum dilution, larval viability is the lowest overall, with 1 and 5% diet concentrations presenting similar but higher viability than 12%. Larval viability is higher at 10^−6^ compared to 10^−4^ inoculum dilution, except for the 12% diet concentration. At 10^−6^ inoculum dilution, no difference in larval viability was found according to diet concentration although the pattern shows a decrease in larval viability as diet concentration increased, as previously observed (FIG 3A and 1A). At 10^−8^ inoculum dilution, larval viability pattern shows an opposite trend compared to 10^−6^, with larval viability increasing with diet concentration. Viability at 1% and 5% diet concentration is similar but lower compare to 12% that displays a significantly higher viability compare to all the conditions tested (FIG 3A). The median larval development time depends on both microbial inoculum and diet concentration although the interaction was not significant (Wald χ^2^ on data transformed by 1/x^2 function, *P*_diet_ = 0.018, *P*_inoculum_ = 1.10e^−14^, *P*_diet x inoculum_ = 0.11). Day_50_ increased upon 10^−4^ and 10^−8^ inoculum dilution compared to 10^−6^, although it varied with the diet concentration. The shortest Day_50_ was measured at 10^−6^ inoculum and 5% diet concentration, being significantly lower compared to 10^−4^ / 12% and 10^−4^ / 1% conditions as well as all 10^−8^ diet concentrations (FIG 3B). Taken together, our results indicate that *Ae. albopictus* larval performance cannot be fully understood without considering the combined impact of diet and bacterial inoculum, that have carry-over effects on adult wing length. While specific larvae survival patterns were observed upon diet x microbiota interaction, the development time seems less constraint by this interaction with a trend for bacteria-associated to develop faster overall especially upon higher microbial inoculum although some diet concentration specific variations still remained.

**FIG 3.**
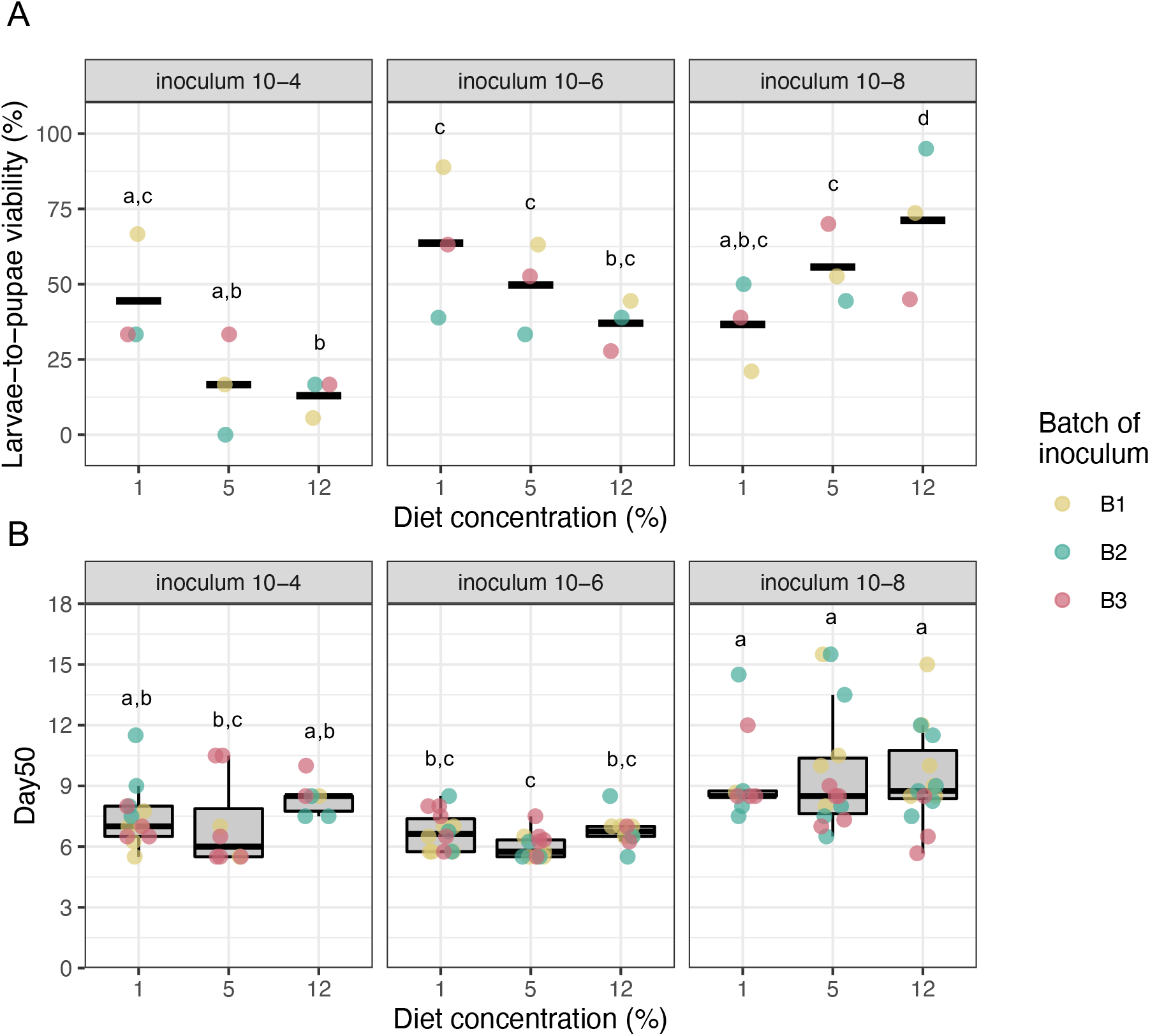
Juvenile performance of *Ae. albopictus* upon concomitant variation of bacterial inoculum and diet concentration. CONV larvae exposed to three microbial inoculum concentrations (10^−4^, 10^−6^ and 10^−8^) issued from three independent batches (B1, B2 and B3, color coded) and three inocula and diet (1, 5, and 12%) concentrations were monitored up to pupal stage. (A) Larval viability expressed as the proportion (in %) of larvae that reached pupal stage at different inoculum and diet concentrations. Each dot represents the mean viability per 6-well plate. (B) Larval development time expressed as the time (in days) needed to reach 50% of the final number of pupae. Each dot represents the mean Day_50_ per well. Different letters indicate statistically significant viability following Tukey-HSD *post-hoc* pairwise comparison. For each batch of inoculum, one 6-well plate was prepared (3 larvae per well) for each inoculum and diet concentration combination.

### Initial bacterial inoculum and diet concentration shape aquatic habitat microbiota composition with bacterial taxa significantly correlated with *Ae. albopictus* performance

Eleven bacterial OTUs were identified at the genus level (with >5% relative abundance) within the rearing water that served as microbial inoculum for CONV larvae (FIG S2). Six out of these 11 OTUs *(Brevundimonas, Delftia, Flavobacteriaceae*, *Pseudomonas*, *Sphingobacteriaceae* and *Sphingobacterium*) were found five days later in the water of CONV larvae (FIG 4A). Visualisations of the relative abundances of the 26 most predominant genera suggested a strong structuration of bacterial microbiota according to initial inoculum concentration, and to a lesser extent, diet concentration (FIG 4A). Beta diversity (microbial diversity between samples) of water samples was represented using non-metric multidimensional scaling (NMDS) with Bray Curtis dissimilarity as the distance metric (FIG 4B-C). When assessing the effect of diet concentration (1, 5 or 12%), inoculum dilution (10^−4^, 10^−6^ or 10^−8^) and batch of inoculum (B1, B2, B3), the triple interaction was significant (PERMANOVA, *P*_inoculum x diet x batch_ = 0.005, R^2^ = 0.0047) (FIG 4B). When trying to disentangle the effect of individual factors on bacterial structure, inoculum dilution had a strong impact when plotted against diet concentration (FIG 4C). These data indicate that complex interactions occur between diet concentration and that the initial composition and abundance of bacterial inoculum shape community structure in larval rearing water. Overall, the bacterial microbiota is homogenous upon when inoculating a high bacterial load whereas a lower amount of inoculum triggers a shift toward a more variable bacterial community in rearing water. Correlation matrix between OTUs abundance (in sequencing reads) and larval traits, including development time (median Day_50_), inoculum dilution (10^−4^, 10^−6^ and 10^−8^), diet concentration (1, 5 and 12%) or larval viability (in %) was performed (FIG 4D). Twenty-two OTUs displayed a significant correlation (Spearman, *P* < 0.05) with a correlation coefficient below −0.4 or above 0.4 that we used as threshold. Only one OTU, assigned to *Bacteroides* genera, correlated with Day_50_ (FIG 4D). Most of the OTUs correlating in relative abundance with one or more larval traits had a low relative abundance with the exception of *Sphingobacterium* (relative abundance correlated with larval viability), *Chryseobacterium* (abundance correlated with both larval viability and inoculum dilution) and *Bosea* (abundance correlated with larval viability and diet concentration) (FIG 4A&D). Details of significant correlation patterns between OTUs relative abundance and larval traits, diet and inoculum concentration for OTUs with the highest relative abundance are shown in FIG S5.

**FIG 4.**
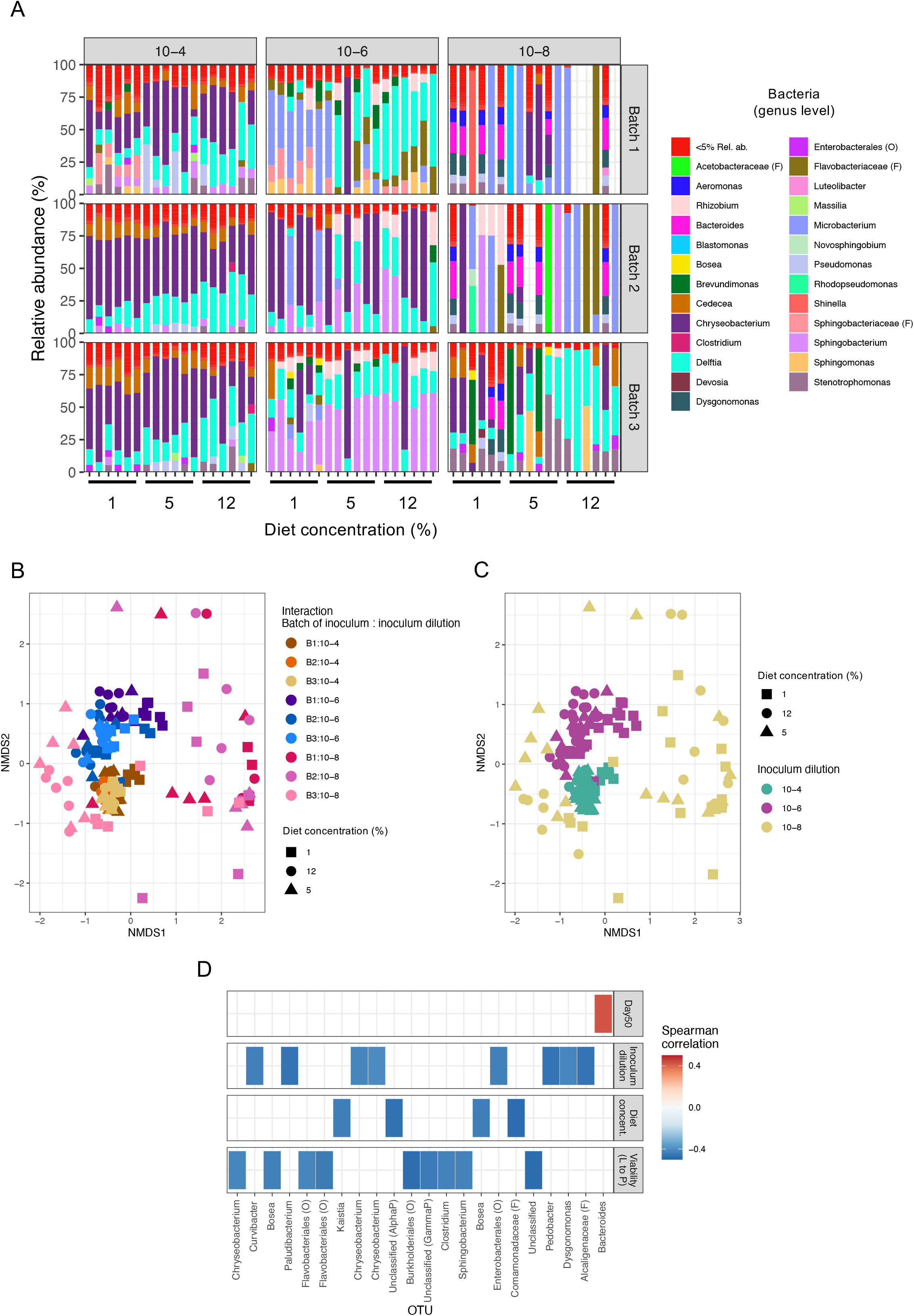
Bacterial community structuration in larval rearing water upon bacterial inoculum and diet concentration gradients. (A) Relative abundances (in %) of bacterial operational taxonomic units (OTUs) at the genus level for each combination of inoculum and diet concentration, for the three batches of microbial inoculum (B1, B2, B3) used. OTUs representing less than 5% in relative abundance were grouped (red, <5% Rel. ab.). (B,C) Non-metric multidimensional scaling (NMDS) analysis of the Bray-Curtis dissimilarities as a function of diet concentration and inoculum dilution (B) or the interaction between batch of inoculum and inoculum dilution (C). (D) Heatmap representing Spearman correlation coefficient between phenotype of interest (development time (Day_50_), inoculum dilution, diet concentration or Larvae-to-pupae viability (L to P)) as a function of OUT abundance. Only OTUs with significative p-value (<1e^−6^) for correlation analysis and a correlation coefficient < −0.4 or > 0.4 are shown. When genus level information was not available, the family (F) or order (O) level is indicated.

## DISCUSSION

Our study demonstrated that microbiota and diet in aquatic larval habitat drive ecological performance of *Ae. albopictus* larvae with carry-over effects on adult mosquitoes. This study is the first, to the best of our knowledge, to address *Ae. albopictus* development using larvae deprived from external microorganisms (AM) and siblings re-associated with conventional bacteria (CONV). AM larvae of *Ae. albopictus* developed up to the adult stage, as observed in axenic from *Aedes aegypti* (36) or *Drosophila melanogaster* (11). If the presence of living but non-cultivable bacteria cannot be ruled out, AM larvae delayed development time and shorter wing length recapitulated developmental phenotypes of axenic *Ae. aegypti* (36). More broadly, these results resonate with studies in axenic flies and mice reinforcing the role of microbiota as a promoter of juvenile development in metazoan (57, 58).

Our results showed that diet and bacteria mostly impact larvae-to-pupae transition. Bacteria from rearing water promoted larval development upon nutrient scarcity but, above a given nutrient concentration, impaired larval viability while the opposite trend was shown in AM larvae. Upon high nutrient concentration, bacterial presence seemed toxic for mosquito larvae although at low diet concentration, it promoted larval development through an increased viability and a shorter development time. Together, it underlies the importance of bacteria and diet interaction on mosquito fitness in *Ae. albopictus.* Bacterial growth in rearing water depends on the strain and diet considered although total bacterial load increases with diet concentration (43, 46). Maintenance of a bacterial homeostasis in mosquitoes is under tight immunological control (65). Therefore, we can hypothesize that diet concentration mediates an increase in water bacterial load that disrupt bacterial homeostasis, leading to a fitness decrease in mosquito.

Despite larval development time depended on diet x microbiota interaction, CONV development time remained around 6 to 7 days with only a small increase at the highest diet concentration tested. This suggests that within a range of diet concentration, bacteria from rearing water can buffer nutrient scarcity that maintains a short larval development time as in *Drosophila* (37). However, development time of CONV mosquito larvae was steadily ~5 days shorter than AM siblings even at the highest diet concentration, conversely to *Drosophila* for which this developmental delay is observed in nutrient-poor conditions but abolished on rich diet (11). Although the use of microorganisms as nutrients by mosquitoes is generally pointed out (66), our results suggest that, beyond a food source, bacteria are actively providing *Ae. albopictus* with compounds as supported by the identification of folate and riboflavin provisioning by bacteria as an essential mechanism for mosquito development in *Ae. aegypti* (41, 42). Further experiments comparing metabolically inactive (*e.g.* heat-killed) and active bacterial exposure in axenic larvae will provide additional evidences to understand host-microbe interactions in mosquito.

A previous study showed that diet concentration impacts microbiota composition in *Ae. aegypti* larvae and adults (46). Our results unravelled a more complex interaction involving both the diet concentration and the bacterial load inoculum in shaping bacterial communities in larval water. Results showed that nutrient-rich water samples colonized with a highly concentrated inoculum have a strongly similar bacterial community compared to nutrient poor samples exposed to a diluted bacterial inoculum. Several OTUs were preferentially retrieved in water following exposure to a highly diluted (10^−8^) inoculum supporting the idea of a more random bacterial community construction upon low initial bacterial dose. Interestingly, the four OTUs that correlated to diet concentration were more abundant at the lowest diet concentration suggesting that nutrient density and/or competition could impact their load upon higher diet concentration. Notably, one OTU from the *Comamonadaceae* family with a higher abundance in rearing water upon low (1%) diet concentration displayed the same abundance/diet concentration pattern in *Ae. aegypti* (46). OTU from genus *Bosea* was more abundant when diet concentration was low (1%), and its abundance correlated with larvae viability although not in a linear way. Present in sylvatic *Ae. aegypti* larval habitats, *Bosea* can trigger a decrease in adult lifespan upon larval exposure (45). *Chryseobacterium* and *Sphingobacterium* abundance negatively correlated with larvae viability. If *Chryseobacterium* was often encountered in mosquito microbiota and restored larval development in mono-association (67), our results suggest that its abundance is key for *Ae. albopictus* performance. *Sphingobacterium* was associated with a developmental delay and shorter wing length in *Ae. aegypti* (43). Our data suggest that it can also impair larval viability in a density-dependent manner. Altogether, our data indicate a complex interplay between bacterial founders and nutrients load that shape bacterial community structure and impact mosquito performance. While it is known that given bacterial strains can compensate for mosquito larvae auxotrophies notably for vitamins (41, 42), data are still lacking to fully understand complex multipartite host-microbe interactions in mosquitoes. For instance, *Lactoplantibacillus plantarum* and *Acetobacter pomorum* triggered a metabolic cooperation that complemented *Drosophila* larval auxotrophies through metabolites that are not produced by bacteria in a mono-association context (68). This underlines that diet-microbiota interaction is key as diet selects for diet-adapted strains, influences the growth of microorganisms and cooperation in the niche which in turn impacts host performance as shown in fly and mammals (4, 5, 69). The situation can be further complexified if we consider other microorganisms from *Ae. albopictus* microbiota that can be directly involved in mosquito metabolism as recently uncovered for fungi (70) or indirect effects such as arbovirus-mediated modulation of bacterial diversity in adult mosquitoes (71).

The development of gnotobiology (76) and holidic diets coupled to in-depth transcriptomic and metabolic analysis will help toward the understanding of mosquito interaction with their biotic and abiotic environment (41). Interestingly, recent work showed opposite conclusions on the impact of microbiota on larval transcriptome, advocating for additional studies that compare transcriptomes of axenic and mono-associated larvae with field-derived bacterial isolates, upon diet and bacterial inoculum variations (39, 40). More broadly, it has potential major applications for alternative vector control strategies for instance by manipulating mosquito oviposition behaviour through bacteria and diet in the rearing water in order to limit mosquito density (75). Altogether, this work provides a better understanding of the ecological determinants of symbiosis in a medically relevant organism while opening interesting avenues for alternative vector control strategies. It also uncovers a fragile equilibrium between diet, bacteria and mosquito fitness, opening questions about the genetic and environmental basis of mosquito traits notably involved in vectorial capacity.

## MATERIAL AND METHODS

### Mosquito colony maintenance

F_9_ and F_10_ from an *Ae. albopictus* colony (referred as VB) established from field mosquitoes collected in 2018 in Villeurbanne and Pierre-Bénite (France) were used. Larvae were maintained at 26°C with dechlorinated water and Tetramin fish food. Adults were raised at 28°C, 80% relative humidity, 16:8 light:dark photoperiod in mass rearing. Females were blood fed on mice in accordance with the Institutional Animal Care and Use Committee from Lyon1 University (Apafis #31807-2021052715018315). Egg papers were stored at 28°C for up to two months.

### Diet plugs preparation

Sterile agar diet plugs of various concentrations were prepared. Finely grinded tropical fish flakes (TetraMin) were mixed with sterile water at final concentrations of 20%, 12%, 10%, 8%, 5%, 2%, 1%, 0.5% and 0.1% (w:v). Bacteriological agar (Conda) was added at a final concentration of 1.6% (w:v). Diet suspensions were autoclaved (120°C, 20 min), poured into 90 mm petri dishes (20 mL per dish) and stored at 4°C for up to 3 days. Die-cut of 0.6 g agar food plugs using a 15-mL sterile tube (Falcon) allowed a precise control of food quantity.

### Preparation of conventional microbial inoculum

Larvae hatched from surface-sterilized eggs were re-associated with a microbiota representative of the insectary-reared siblings to obtain conventional (CONV) larvae. Briefly, VB eggs hatched were in hatched for 2h at −20 ATM in a vacuum chamber then 200 first instar larvae were transferred in 24 x 32 x 9 cm plastic trays with 1.5 L of dechlorinated water. After 7 days at 26°C with 0.1 g / tray every two days of grinded fish flakes (Tetramin) supplemented with yeast extract (Biover) (3:1 w:w), 50 mL of water (without larvae) from 3 independent trays were pooled to constitute the CONV inoculum. When testing for inoculum batch effect, 3 batches were prepared from independent egg papers. To prepare CONV larvae first instars from surface-sterilized eggs were exposed immediately upon hatching to CONV inoculum at selected dilution (from 10^−3^ to 10^−8^) in sterile water (Gibco). Bacterial composition of CONV inoculum is provided (FIG S2) while no eukaryotic microorganisms were detected by 18S PCR.

### Conventional and altered microbiota mosquito production

After hatching larvae acquire their microbiota through the ingestion of microorganisms present on the egg surface or in the rearing water. *Ae*. *albopictus* larvae with altered microbiota (AM) were obtained by surface sterilization of mosquito eggs (FIG S1). In 6-well plates, a 0.6 g agar food plug at the selected concentration was added per well with 5 mL of sterile water (for AM condition) or 5 ml of CONV inoculum at selected dilutions (for CONV condition). A total of 3 larvae / well was added and plates were incubated at 28°C in complete darkness for up to 22 days. The well is the biological replicate unit and 6 to 18 wells were tested for each condition. For each experiment, agar plugs and PBS (in which surface-sterilized larvae hatched) were incubated on LBm agar for 7 days at 30°C to control for the absence of cultivable contaminants. Wells were observed daily and discarded in presence of turbid water. As a majority of microorganisms are not cultivable, random wells were observed for each plate under the microscope to assess the absence of microbial contaminants.

### Larvae viability and development time

Presence of pupae was recorded daily at fixed hours. Larvae-to-pupae viability represents the percentage of larvae that reached pupal stage. Development time represents the time (in days) needed to reach 50% of the total number of pupae (Day_50_). On the day of emergence, pupae were individually transferred with ~300 μL of rearing water in a sterile tube for emergence. Adults were sexed before storage together with water of emergence at −80°C.

### Wing length measurement

Wing length is a proxy for *Ae. albopictus* adult performance (79). Adults stored at −80°C were thawed and both wings were dissected under a Leica M80 stereomicroscope. Wings were included Eurapal (Roth) on a 10-wells, epoxy-coated glass slide (Labelians). Slides were photographed at 20x magnification with Leica MC170 HD camera. Images were analysed with ImageJ (version 2.1.0/1.53c). Wing length was measured between the intersection of the second and third vein and the intersection of the seventh vein with the wing border using ImageJ (80).

### DNA isolation from larvae rearing water

Five days post exposure with CONV inoculum, 500 μL of rearing water were stored at −20°C prior to total DNA isolation. Samples were centrifuged for 20 min, 4°C, at 17,000 G and the pellet containing the microorganisms was used to extract DNA with the DNeasy Blood & Tissue kit as recommended (Qiagen). DNA concentration was estimated by Qubit dsDNA HS kit (Thermo Fisher scientific) and samples were stored at −20°C. A blank control was performed by using only DNA lysis buffer to control DNA microbial contaminations arising from kit reagents or the environment during DNA isolation.

### Quantitative PCR of 16S DNA

The 16S DNA load was measured by qPCR using the Itaq SYBR green supermix kit (Bio-Rad), 784F (5’-AGGATTAGATACCCTGGTA-3’) and 1061R (5’-CRRCACGAGCTGAC’) primers and 5 ng of template DNA isolated from water samples. The 16 μL reaction comprised 0.48 μL of each primer at 10 μM, 8 μL of Master mix and 5.04 μL of PCR grade water. After a single denaturation step at 95°C for 3 min, a two-step amplification was performed including 10 s at 95°C followed by 30 s at 60°C, for 40 cycles on a Bio-Rad CFX96 machine. The number of 16S copies per μL was calculated using serial dilutions of an *Acinetobacter* PCR amplicon from 10^8^ to 10 copies/μL.

### Microbial amplicon sequencing

Identification of bacteria and eukaryotes was based on PCR amplification of a ~280 bp fragment of the V5-V6 variable region from the 16S rRNA gene (52) and a ~430 bp fragment of the 18S gene (81). Duplicate PCR for each sample were done using the 5X Hot BIOAmp (Biofidal, Vaulx-en-Velin, France – http://www.biofidal.com). Duplicates PCR products were pooled and 5 μL were separated by electrophoresis on 1.5% agarose gel supplemented with 2.5 μL of clear sight DNA stain for 17 min at 100V. All the 18S PCR were negative. For the 16S, all PCR were positive at expected size. A total of 188 libraries from 16S amplicons were constructed, including controls. Sequencing was done on Illumina MiSeq (2×300 bp, paired-end) at Biofidal. In total, 20,764,855 reads were obtained and demultiplexed. Sequence quality control and analysis were carried out using the FROGS pipeline (82) as described (83). Taxonomic affiliation was performed with SILVA database 138.1 for bacteria (84) with Mothur pipeline (85) at a 80% minimum bootstrap using a naïve Bayesian classifier (86). Sequences were grouped into operational taxonomic units (OTUs) by clustering at 97% similarity. No OTUs were detected in the blank extraction or PCR controls. To compare samples, normalization was performed at 11,780 sequences. A total of 184 operational taxonomic units (OTUs) was obtained. OTUs with a relative abundance less than 10 times greater than that observed in the negative control were removed (52). All FastQ files were deposited in the EMBL European Nucleotide Archive (https://www.ebi.ac.uk/ena) under the project accession number PRJEB57586. The small amount of residual bacterial DNA detected in water from altered microbiota larvae was quantified by qPCR, and taxonomic identity and relative abundance of the 5 OTUs detected shown (FIG S3).

Statistical analysis. Analyses and graphical representations were done on R (http://www.r-project.org/). Larvae- and pupae-to-adult viability (binary response variable) were analysed by generalized linear mixed-effects (GLMM) (87). GLMM with a binomial distribution and a probit link function were fitted by maximum likelihood (Laplace approximation). The development time (Day_50_) and wing length (in mm) were analysed using linear mixed-effects (LMM) models fit by restricted maximum likelihood while controlling for normal distribution of the residuals. Microbial status and diet concentration in interaction represented fixed effects while experiment or batch of inoculum were included as random effect. The GLMM and LMM models were conducted under *lme4* package version 1.1-25 (88). The inference of explanatory variable on variations of the response variables was tested with a Wald X^2^ test or an ANOVA for binomial and continuous data respectively. *Post-hoc* comparisons after GLMM/LMM were performed with *emmeans* package version 1.5.2-1 (89) to estimate pairwise differences between diet concentrations and microbial status using Tukey-HSD test with p-value correction for multiple comparisons. Adonis-ANOVA and Non-Metric Multidimensional Scaling ordination were performed with the ade4 and vegan packages (90, 91). Non-linear Spearman correlations were performed with the Hmisc package (92). Other R packages were used for data organization and representation such as *plyr* version 1.8.6 (93) and *ggplot2* version 3.3.2 (94).

## ACKNOWLEDGMENTS

This project was funded by IDEX Lyon scientific breakthrough project, Micro-Be-Have. We acknowledge the contribution of the DTAMB and ACSED platforms of the Research Federation BioEEnviS (Biodiversité, Eau, Environnement, Ville et Santé) supported by University Lyon1 and CNRS. We thank all the members from the Micro-Be-Have consortium and the G-RREMI (Groupe Régional de Recherche En Microbiologie des Interactions Auvergne-Rhône-Alpes) for insightful comments and discussions.

---

**FIG S1** Protocol for surface sterilization of *Ae. albopictus* eggs. Protocol was adapted from previous work (36) *Ae. albopictus* eggs (<2 months old) were used. Careful visual inspection of egg papers under binocular was conducted prior to each experiment to avoid concave eggs (as a proxy for unfertilized eggs) or egg papers with mold or mites. Within each experiment, at least three different batches of embryos from the same generation but laid at different days were used. Under safety cabinet, eggs were rinsed twice by dipping the paper in sterile water to discard debris. Egg papers were then soaked in a petri dish containing 70% ethanol solution (in sterile water) for 5 min then transferred in a 50-mL falcon tube containing 30 mL of sodium hypochlorite (3% active chlorite) supplemented with 4 mg/mL ampicillin for 5 min. Sodium hypochlorite immediately detached the eggs from the paper, that was removed using clean forceps. Within the 5 min, eggs quickly settled at the bottom of the tube allowing the complete removal of sodium hypochlorite without the need of centrifugation. Eggs were resuspended in 25 ml of 70% ethanol and incubated for 5 min. We ensured that all eggs remained immersed in ethanol and gently agitated the tube to allow a complete contact of the eggs with ethanol solution. Eggs were rinsed three times in sterile water (Gibco) for 5 min then 30 mL of sterile 1X PBS was added. The tube was closed with a 0.2 μm filtered cap and transferred in a vacuum chamber outside the cabinet for 40 min at −20 ATM to allow hatching. The sterile first instar larvae were transferred under the safety cabinet in a sterile petri dish and immediately transferred in 6-well plates using a P1000 pipette according to the experimental design.

**FIG S2 Taxonomic identity and** relative abundance of bacterial OTUs in water inoculum for CONV larvae. Relative abundance (in %) of operational taxonomic units (OUT) of bacteria at genus level (or family (F) level) in water of conventionally reared larvae that served as inoculum for CONV condition. Two DNA samples (S1, S2) per batch of inoculum (Batch 1, 2 and 3) were sequenced except for Batch 3 (only one sample). The two OTUs that were isolated by culture-dependent approach on LBm media are shown (*). OTUs representing less than 5% in relative abundance were grouped (red, <5% Rel. ab.).

**FIG S3** Bacteria composition and relative abundance of altered microbiota larvae rearing water DNA samples. (A) Quantification of 16S copies number (log_10_) in DNA isolated from water of conventional (CONV) and altered microbiota water samples after five days at 28°C. CONV samples (Cq values from 14 to 16, mean 5.5×10^6^ copies) originated from water containing larvae exposed to 10^−8^ dilution of inoculum and incubated with 5% diet concentration. The altered microbiota DNA samples (Cq values from 28 to 29.4, mean 6.39×10^2^ copies) originated from water with diet but no larvae (S1, S2), water with larvae and diet (S4, S5) or water without larvae nor diet (S3). A blank DNA isolation control (Cq 34.9, 10.5 copies) was done by replacing water with DNA template by lysis buffer (S6). (B) Relative abundance of bacterial operational taxonomic units (OUT) at genus level in AM DNA water samples (S1 to S5). OUT representing less than 5% in relative abundance were grouped (red, <1% Rel. ab.). No reads were obtained from the blank DNA control. Presence/absence of larvae and diet is indicated for the five samples.

**FIG S4** *Ae. albopictus* larval viability depends on microbial exposure and diet concentration. Larval viability expressed as the proportion (in %) of conventional (CONV) exposed to a low dilution (10^−3^) of microbial inoculum (A) or altered microbiota (AM) (B) larvae that reached pupal stage upon different diet concentrations (in %). Each dot represents the mean viability of 3 wells from two independent 6-well plates (3 larvae/well).

**FIG S5** Correlation patterns between relative abundance of selected OTUs and larval traits or experimental factors. Only the most abundant OTUs for which a significant correlation (Spearman, *P* < 0.05) with a correlation coefficient below −0.4 or above 0.4 was found are displayed.

